## Supplementary figures and images for "Meta-Analysis of Oxidative Transcriptomes in Insects"

### FigureS1

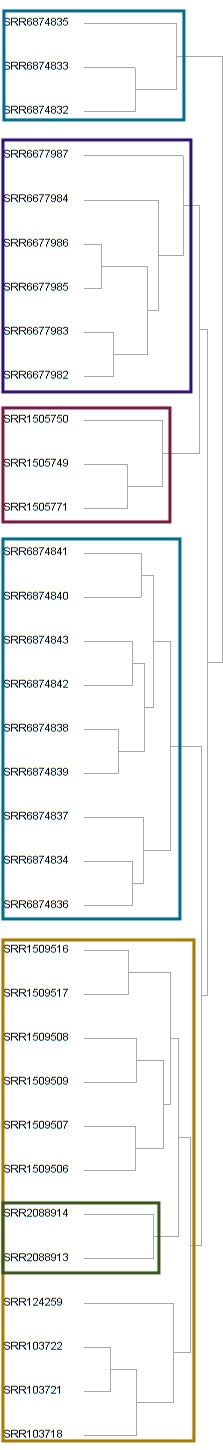
